# Taking a landscape approach to conservation goals: designing multi-objective landscapes

**DOI:** 10.1101/2020.01.21.914721

**Authors:** Anna R. Renwick, Alienor L.M. Chauvenet, Hugh P. Possingham, Vanessa M. Adams, Jennifer McGowan, Vesna Gagic, Nancy A. Schellhorn

**Affiliations:** School fo Biological Sciences, University of Queensland, Brisbane, Queensland 4072, Australia.; Environmental Futures Research Institute, School of Environment and Science, Griffith University, Gold Coast, QLD, Australia; The Nature Conservancy, 4245 North Fairfax Drive, Suite 100, Arlington, Virginia 22203, USA.; Discipline of Geography and Spatial Sciences, University of Tasmania, Hobart, Tasmania 7000, Australia; CSIRO, Ecoscience Precinct, GPO Box 2583, Brisbane, Queensland 4001, Australia.

**Keywords:** Land sharing land sparing, Decision Theory, Agriculture, Biodiversity, Forestry, Urban, Habitat conversion, Management objectives

## Abstract

Designing landscapes to accommodate both humans and nature poses huge challenges, but is increasingly recognised as an essential component of conservation and land management. The land-sparing land-sharing framework has been proposed as a tool to address this challenge. However, it has been largely criticised for its simplicity. We provide a new conceptual framework amenable to the application of structured decision-making that moves beyond the dichotomy of land-sparing or land-sharing. Using this new framework, we present a general system model that can be used to make land management decisions for the conservation of species, ecosystem services and production land at different spatial scales. The model can be parameterised for specific systems using information about: the current state of the landscape, the rates of change between landscape states, and the cost and effectiveness of taking actions. To demonstrate the utility of the model we apply it to three different landscape types. Across our three case studies, we show that investment into one of three management actions (varying degrees of management and restoration) can move the system towards more biodiversity or more managed land depending on the objectives of the land manager. We show that the dynamic and flexible nature of the landscape is important to take into account rather than a static snapshot in time. Rather than focusing on establishing the perfect landscape with a set proportion dedicated to production and to biodiversity conservation, we argue that a more useful approach is to establish incremental movements towards a landscape that meets the goals of multiple objectives. Our framework can be used to illustrate to decision makers the costs and trade-offs of different actions and help them determine land management policy.

## Introduction

Globally, nations have committed to halt further extinctions and safeguard biodiversity (Convention on Biological Diversity 2011b). The continued expansion of protected areas (PAs) and effective management of these areas are central components of achieving this goal; the Convention on Biological Diversity Aichi Target 11 commits signatory nations to protect 17% of terrestrial and 10% of marine environments by 2020 (Convention on Biological Diversity 2011a). Achieving this target requires taking actions to ensure that there is an adequate amount of intact land within the landscape. In addition to commitments to maintain intact land, there are global and regional level policies to restore degraded ecosystems; for example, Aichi Target 15 commits nations to restore 15% of degraded ecosystems by 2020 (Convention on Biological Diversity 2011a). However, there is the often conflicting requirement to have sufficient areas to support human needs (i.e. agriculture production, forestry, urban areas) (Niemelä et al. 2005; McDonald et al. 2008; Tanentzap et al. 2015), and hence achieving these goals are not without great challenges.

Determining the optimum design of landscapes to accommodate both nature and human land use has become a hot topic in the conservation world (Goulart et al. 2016). This is of no surprise given the rapid increase in human population size (Cincotta et al. 2000) and inevitable encroachment into areas once solely reserved for nature. Land sharing or land sparing is one conceptual approach to thinking about balancing the needs of nature and humans – should we design landscapes with a focus on intact protected natural areas (spared) or where benefits to humans and nature are met in the same place (shared)?

The land sparing versus land sharing framework emerged from the Borlaug hypothesis that claimed that the Green Revolution of intensification of agriculture had ‘spared’ several hundred millions of hectares of land that would have been converted into agriculture. It was further developed by Green et al. (2005) and Balmford and Bond (2005), primarily to determine the optimal strategy for conserving species while maximising yield in agricultural landscapes. In the agriculture context, land sparing involves segregating the landscape by intensifying agricultural production, thereby reducing the area required to meet production demands in one part of the landscape, while leaving the remaining land for nature. This has the expectation that increasing intensification will increase yield and reduce the area required to meet production demand (Phalan et al. 2011). In contrast, land sharing involves integrating agriculture and biodiversity preservation together, or in close proximity, through the use of less intensive production methods over a larger area to meet the same production levels (Phalan et al. 2011). The framework has caused quite a divide in the ecological sector with strong advocates for and against. One of the main falsities by its critics has been its oversimplification, both in terms of the dichotomy approach and also in its implementation (Kremen 2015).

The framework is now being used in a broader context to encompass issues about landscape design in general, and not just specifically about maximising yield, for example the design of cities and plantation landscapes, livestock production, and fisheries production (e.g. Lin & Fuller 2013; Edwards et al. 2014; Law et al. 2016; McGowan et al. 2018). As its application moves into more general usage the issue of oversimplification becomes even more apparent, emphasizing that limiting landscape decisions to two states (shared or spared) is often too simplistic to produce viable landscape design options. Managing landscapes that balance the needs of nature and humans requires consideration of the myriad of land use options between these extremes. We argue that landscape scale decisions are complex and dynamic, and that land management decisions will likely require consideration of multiple objectives.

We provide a new conceptual approach to landscape design grounded in a system model for landscape dynamics and structured decision-making. Our approach proposes a path beyond the, often false, dichotomy of either sparing or sharing. Rather than aiming for a utopian end- point landscape implied by the land sharing land sparing framework, we aim for small incremental changes to the system, which result from investing in various management actions. This reflects a more realistic view of plan implementation that is often constructed of incremental system changes to move towards a preferred end point (Pressey et al. 2013). Our general system model provides a visual tool to conceptualise and illustrate the relative balance of land use, and the direction the landscape is moving in under the current management regime, both of which can provide information to aid in developing high level policy guidelines. The model can be used to make the best decisions that balance benefits to any variety of variables, for example the conservation of species, ecosystem services and production, at different spatial scales. To demonstrate the framework’s utility, we apply it to three different landscape types: urban, forestry and agriculture.

## Modelling framework

We develop a generalizable model that can be used to better understand when and where to implement different conservation actions while accounting for their costs and benefits at the landscape level. The aim of a system model is to understand the management options that are possible for the system, and make the trade-offs between decisions explicit. Our model accounts for the current state of the system (e.g. amount of land protected or converted), the rates of change between the different states caused by anthropogenic threats or as the result of conservation actions (and their interactions), and the cost and benefits of taking actions. The best course of action will be dictated by the current state of the system (i.e. proportion of land protected or converted) and the overall objective for the landscape. This qualitative approach allows the model to be applied to landscapes at any spatial scale, overcoming one of the issues highlighted in the land sparing land sharing framework whereby an area may be classified as land sparing at the farm level but land sharing at the regional level (Fischer et al. 2014). The focus of this model is not to define a utopian landscape or to replace the usage of the spatially explicit models such as Marxan and Invest used in conservation planning, but instead to indicate management pathways for incremental changes to move the system towards a more desirable state based upon the multiple objectives of the landowner.

### System Model

We first define a landscape with three broad but common land use states (these can be applied to various systems and are not restricted to agroecosystems): Converted habitat for human use (C), Intact habitat (I), and Modified habitat which is a mix of natural and human use (M) (see Box 1 for definitions). We assume these different states deliver varying benefits for biodiversity. For example, a fully converted habitat state (C) tends to have a low to negligible benefit in terms of enhancing biodiversity, while the intact habitat state (I; here encompassing both formally protected and unprotected areas) delivers the most benefit to biodiversity. Modified habitat (M) presents moderate (and potentially different) benefits to biodiversity, which vary depending on how sensitive a species is to a particular threat and how modified (different from intact) the habitat is, and can be beneficial for the provision of ecosystem services such as pollination and natural pest control (Wright et al. 2012). *(e.g. C< M< I*)

##### Box 1 List of system state variables

###### Converted (C)

Habitat that is cleared and actively used for human uses (e.g. housing, agriculture, intensive monoculture forestry).

###### Intact (I)

Habitat that is intact (i.e. not converted and not degraded by other environmental threats) and thus maintains its native species and ecosystem functions. It may be formally protected or not.

###### Modified (M)

Habitat that is at least partially cleared or modified for mixed land uses but retains some of its native species and ecosystem functions, as well as production-based ecosystem services. For example, grassy strips within cereal crops or parks within cities.

Our non-spatial, three state model offers a simplistic basis that can be coupled to complex benefit functions that incorporate variable benefits associated with different system states, for example food production on modified or converted lands, and biodiversity on modified or intact lands. The benefits of this modelling approach is that it can account for benefits across different land uses. Depending on the managers’ objectives for a landscape, and given the relative contributions of different states to biodiversity outcomes and production values, management actions are needed to shift habitat between the three states. This requires understanding not only the system’s current state and desired usage, but also the relative rates of loss of habitat through active conversion for production and from other threatening processes, as well as the conservation actions that can be implemented to address these threats (Fig. 1a). Here, we present a simplified model where habitat loss stems from two processes (*δ_i_*): active conversion (δ_c_) which moves both intact (I) and modified (M) land towards converted land (C), and habitat modification (δ_m_), which moves intact land (I) towards a modified state (M). Management actions, such as intensive restoration (b_1_) move C to I, moderate restoration (b_3_) move M to I, and land management such as land management (b_2_) move C to M.

**Figure 1:**
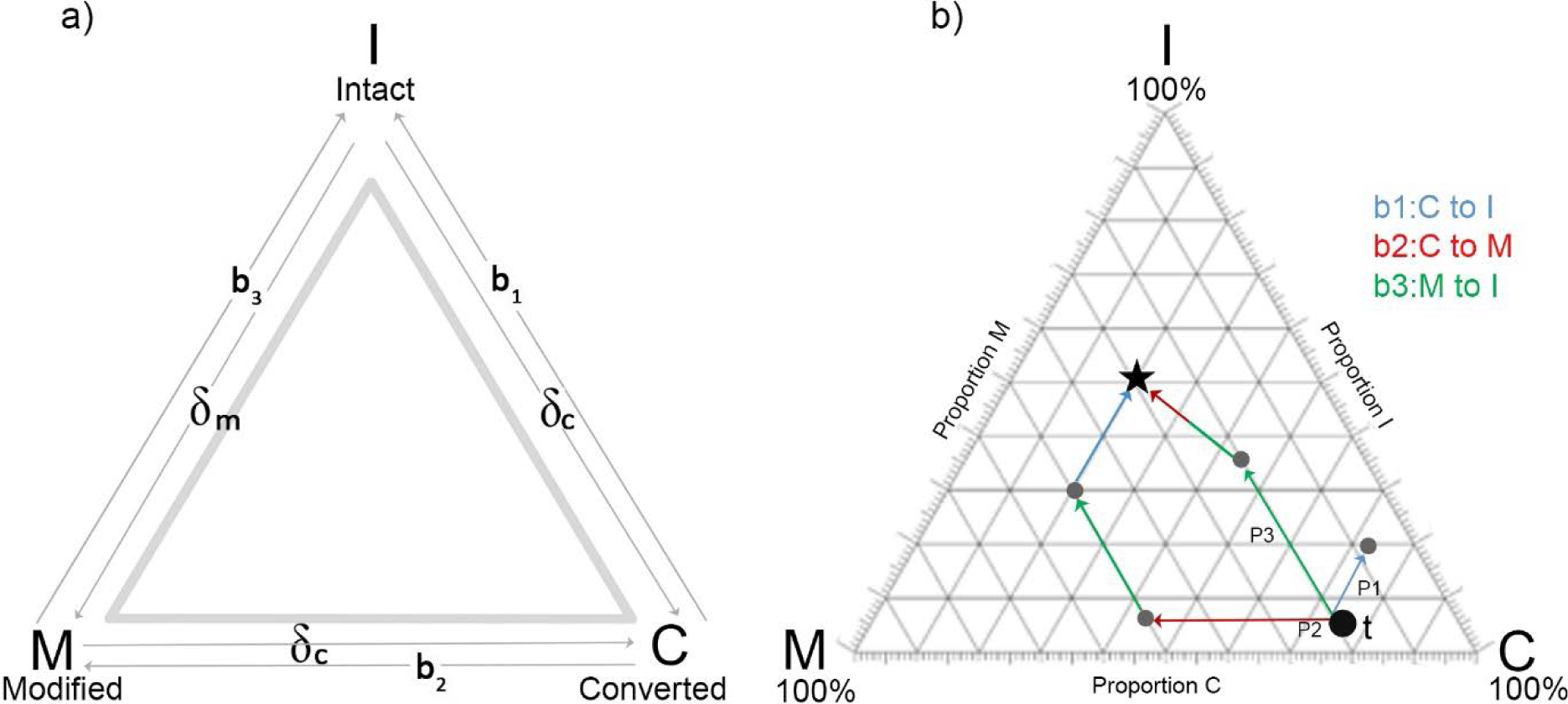
Three state system model. a) System state model indicating rates of loss due to threat (δ_i_) and conservation management actions (b_i_), b) A hypothetical example to illustrate our framework. In order to move from the current state at time, *t* (20% Modified Areas, 75% Converted Areas, and 5% Intact Areas) towards a desired landscape (30% Modified Areas, 20% Converted Areas and 50% Intact Areas; illustrated by the star), a manager can choose from a variety of strategies (P1-P3) depending on the planning horizon and preferred actions b_1_ (blue), b_2_ (red), or b_3_ (green).

The model (Fig. 1a) is described by the following set of differential equations:

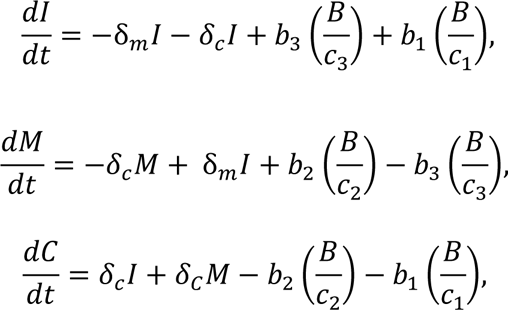

where *δ_c_* is the rate of land conversion while *δ_m_* is the rate of land modification. The parameters, *b*_1_, *b*_2_ and *b*_3_, are the percentage of the total budget (*B*) spent on the three conservation interventions (intensive restoration, land management and moderate restoration), such that *b*_1_ + *b*_2_ + *b*_3_ = 100%. Each conservation action *i* has a different associated cost per unit area (*c_i_*).

We illustrate how to operationalize our framework first with a hypothetical example (Fig. 1b). Imagine the current state of the system at time *t*, consists of 20% Modified Areas, 75% Converted Areas, and 5% Intact areas. The desired landscape consists of 30% Modified areas, 20% Converted Areas and 50% Intact Areas. The optimal action can change at each time step based on the state of the system. There are many pathways a manager may choose to achieve the desired landscape. The decision about which action or set of actions to invest in can vary depending on the planning horizons, objectives, budgets and the feasibility of the actions. For example, a manager may only be interested in the optimal strategy in the near-term (one time step) that gets them closest to the desired state (Fig. 1b, Path P_1_). In our example, this strategy leads to an investment in action b_3_, resulting in a landscape that is now 20% M, 60% C, and 20% I. Another approach could be to optimize over a much longer planning horizon (i.e. several decades into the future) to fully realize the desired state (Fig. 1b, Path P_2_ or P_3_).

In some cases, the best strategy may be to first allocate the entire budget in one action, but then to split the allocation between two actions simultaneously in the next time step (as illustrated in Fig.1b, Path P_2_). In another instance, sequential investment in different actions may be the optimal long-term strategy (Fig.1b, Path P_3_). How a manager chooses the optimal strategy can vary according to the context but could include achieving the desired state in the most cost-effective way possible, the fastest way possible, and/or where the transitioning landscapes at each time step are all acceptable compositions of M, C, and I, that are better than the state at time *t*.

We further illustrate our model using three case studies from different landscapes (urban, forestry, agriculture), each with multiple objectives (see Fig. 2 for mapping of case studies to framework with initial conditions and considered pathways of change). For each case study we show the different pathways that can be taken to achieve the desired landscape state taking into consideration the landownes multiple objectives. We also look at the effect of overall budget in achieving the multiple objectives running scenarionswhere the costs of the different management actions are all equal (c1=c2=c3) but the overall budget is a) small, b) medium and c) large.

**Figure 2:**
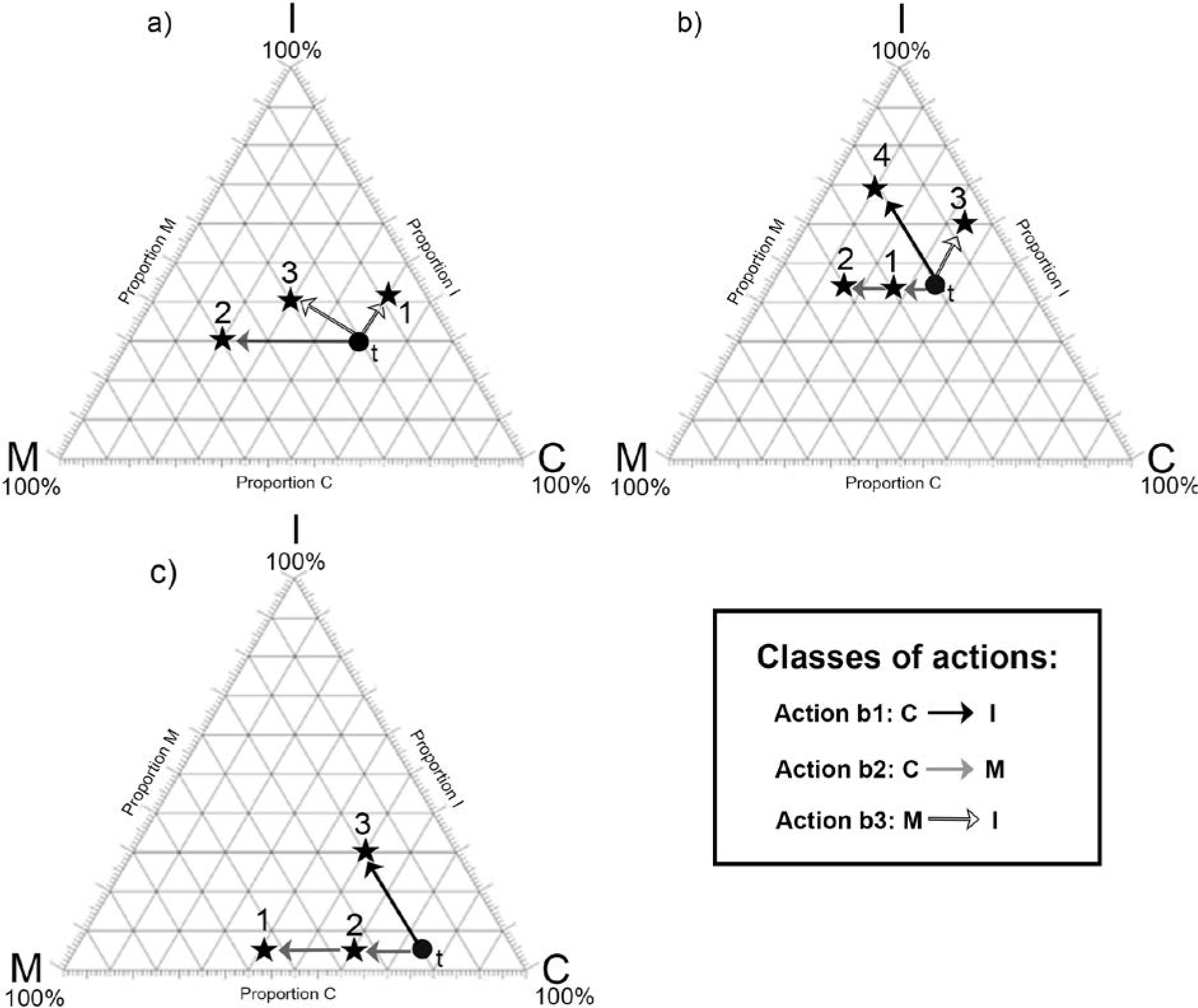
Three case study applications of system model. A) urban application with initial condition (t) of I=30%, M=20% and C=50%. We consider three potential pathways: 1) return cleared habitat to intact habitat (C to I); 2) increase the intensity of dwellings in an area to allow for a greater area to be designated as parkland (C to M); 3) increase the amount of I habitat through returning M to I. B) Forestry application with initial condition (t) of I=44%, M=20%, C=36%. We consider four potential pathways: 1) modify management and establish plantations on previously cleared sites (C to M), 2) plant fast growing timber trees on previously cleared sites or clear new sites (C to M), 3) allow forest land to be used for other non-timber activities e.g. carbon storage instead of being cleared (C to M)), or 4) protect and retain old growth primary forest while also restoring cleared land to an intact forest state (C to I). C) Agriculatural application with initial condition (t) of I=5%, M=20%, C=75%. We consider three potential pathways: 1) restore cleared habitat to accommodate both agriculture and biodiversity to increase yield and farm income while minimizing the risk to biodiversity (C to M), 2) manage habitat for natural pest control on their land (C to M), 3) take land out of production and revegetate, or improve non-productive land (C to M, C to I or M to I).

## Case studies

Each of the three case studies have multiple objectives, actions, costs and constraints based on the desires of the landowner (Fig. 2). Our case studies are hypothetical and different variables can be used depending on the landscape the model is applied to.

### 1. Urban

#### City planning and biodiversity

Urban areas are the most human modified ecosystems on the planet and are globally expanding faster than any other type of land use (Seto et al. 2011). Cities are often built on previously highly productive ecosystems with rich biodiversity (Imhoff et al. 2004).

Consequently, urbanisation is considered a highly threatening process for biodiversity and food production, and given the speed and extent of urbanisation, reconciling biodiversity with urban development could not be more imperative. The land sparing versus land sharing framework has been proposed as a mechanism to try to address this challenge (Lin & Fuller 2013). Theoretically, land sharing cities consist of low density cities with small green spaces such as small parks and private gardens but lack large parks or reserves. Conversely, land sparing cities consist of high density built-up areas in a smaller area allowing large green spaces in the form of large contiguous parks or reserves to be set aside for nature, such as London or New York (Lin & Fuller 2013; Stott et al. 2015). However as in the agricultural context, this dichotomy is an unrealistic description of the world’s cities and a continuum in intensity is more realistic especially as urban intensity is rarely uniform and varies within individual cities themselves (Stott et al. 2015).

An urban landscape can broadly be divided into three states: 1) I, reserve (place largely made up of native vegetation that is primarily for conservation and nature-based human amenity outcomes), 2) M, small park or unallocated land (a place that is not houses, industry or transport but is urban greenspace used for a wide variety of human cantered purposes not primarily nature), and 3) C, houses and other infrastructure (a built-up area with no provisions for nature). A typical overarching objective for city planners is to optimise housing and amenities while still provisioning green spaces that benefit both humans and biodiversity.

#### Applying the decision framework: Brisbane, Australia case study

We use Brisbane, in the subtropical region of Queensland, Australia as an example to illustrate the use of the framework for urban planners to maximise biodiversity while accommodating the increasing housing demand and human needs. Brisbane’s new city plan aims to restore 40% of the city area to natural habitat by 2026 (Brisbane City Council 2014). However, it is also at the centre of the fastest growing regions in Australia with an inner city predicted population to grow by c.28% by 2031 (Brisbane City Council 2012). Consequently, it is estimated that the city will need approximately 156,000 additional dwellings by 2031.

##### a- Objectives

Urban planners have objectives to: 1) maximise the quality and quantity of housing to meet the upcoming demands within the budget available and without removing any intact habitat (i.e. increase/maintain C or M without decreasing I), 2) provide access to parks and informal open space for people (e.g. green space within 400m walk or five minute drive for inner city residents; Brisbane City Council 2012; i.e. increase or maintain M), and 3) retain a certain amount of biodiversity and examples of the original ecosystems for cultural reasons and to avoid/minimise local extinctions (Brisbane City Council 2017; maintain I).

##### b- Actions

In order to achieve the objective of increasing housing, amenities and revenue stream to accommodate the rising population and requirement for 156,000 additional dwelling while restoring 40% of city area to natural habitat, there are a number of actions that planners need to consider:1) modified management to increase the area of managed parkland to ensure every house has easy access to green space by replacing low rise suburbs with high rise dwelling and more parkland (C converted into M), in addition this may also increase with amount of housing available without altering the amount of habitat in each state, and 2) land management to increase the amount of native vegetation to achieve the 40% natural habitat goal by buying and revegetating parkland and bushland with native species, and partnering with residents to restore privately owned bushland (M converted into I).

##### c- Costs and constraints

Meeting the infrastructure demand of the growing population while achieving this 40% natural habitat goal will require innovative development plans including smaller apartments, high rises, and use of cleared but empty sites. Determining the optimum development plan is further complicated by the constraints given to each action and budget limitations, and will therefore involve trade-offs that need to be carefully assessed in terms of costs. A further constraint to the model is that reserve land should not be developed (I changes to C) and it is unlikely that intensively used land will be returned to reserve (C changes to I).

##### d- Model

We estimated values of I=30%, M=20% and C=50%, and δ_c_ (rate of land conversion from state M to state C) = 0.6% while δ_m_ (rate of land modification from state I to state M) = 0.0%, based on the aims and objectives of the Brisbane City Council (Brisbane City Council 2012). This means that under the current conditions both M and I are being converted to C. As cleared habitat (C) is very unlikely to be converted to intact habitat (I) we set investment into b_1_ at 0% which transforms the problem into a two state problem.

We show three pathways from the initial starting point that a manager many choose to invest their efforts in order to achive their objectives (Fig. 2a). The first pathway is to return cleared habitat to intact habitat. This is very unlikely pathway to occur but is shown for visual purposes. The second pathway is to increase increase the intensity of dwellings in an area to allow for a greater area to be designated as parkland (C to M). The third pathway is to increase the amount of I habitat through returning M to I. Regardless of the budget size, we found that in order to achieve the second pathway of assuring every house has access to green space (increasing M), the majority of financial investment has to be into land management (b_2_) (Supplementaty figure 1A). In order to achieve the third pathway of increasing the amount of green space available and retaining examples of original biodiversity and ecosystems, the amount of I habitat needs to increase. To maximise the amount of I, the majority of investment should go into moderate restoration (b_3_) (Supplementary figure 1)).

### 2. Forestry

#### Forestry practices and biodiversity

Deforestation and forest degradation are well known drivers of biodiversity loss globally. Timber harvesting including clear-felling and selective logging, plantations, fire, and fragmentation can all negatively affect forest biodiversity resulting in localised extinctions and changes in community composition (Meijaard et al. 2005; Edwards et al. 2014). Forest managers have a challenge to reduce threats to biodiversity and limit carbon emissions from forest degradation while maintaining long term timber supplies and economic viability (Putz et al. 2012).

Under the land sparing versus land sharing framework forests can be intensively logged within part of the concession while leaving the rest of the area unharvested (land sparing). Alternatively the forest can be harvested at low intensity across the whole concession (land sharing) (Edwards et al. 2014). Our approach goes beyond this simplistic dichotomy and divides forests into three types of land use 1) intact primary forest (I); 2) disturbed or second growth forest (M); and 3) cleared areas (C).

#### Applying the decision framework: Borneo case study

Borneo has lost forest cover at almost double the rate of the rest of the world’s humid tropical forests (Achard et al. 2002). Consequently, approximately 34% of old growth forest has been cleared between 1973-2015 (Gaveau et al. 2016). Protecting forests from conversion to plantations, fire and illegal logging is crucial to reducing deforestation rate in Borneo.

##### a- Objectives

Forest managers have a number of objectives to meet in relation to increasing the amount of modified and/or intact habitat: 1) conversion of cleared areas to plantations for economic benefits (i.e. decrease C and increase M), 2) carbon storage (i.e. maintaining or increaseing M or I), and 3) biodiversity conservation (i.e. increase I). The latter two objectives are relatively new within forest management but are gaining recognition for the benefits, including economic, both directly and indirectly (Bekessy & Wintle 2008; Wells et al. 2013).

##### b- Actions

In order to meet these objectives, forest managers can: 1) modify management and establish plantations on previously cleared sites (C to M), 2) plant fast growing timber trees on previously cleared sites or clear new sites (C to M), 3) allow forest land to be used for other non-timber activities e.g. carbon storage instead of being cleared (C to M)),or 4) protect and retain old growth primary forest (C to I) (Fig. 2b).

##### c- Costs and constrains

Retaining natural forest and minimising their conversion to other uses including plantations is priority. However, plantations bring high short term economic benefits to the country and are a substantial component of the country’s economy (Fisher et al. 2011). Political decentralisation and instability, and lack of clear laws governing forest lands constrain the ability to protect and preserve old growth forest (Gaveau et al. 2016).

##### d- Model

In a recent study, Gaveau et al. (2016) estimated the forest and industrial plantation areas in Borneo as I=44%, M=20%, C=36% and δ_c_ (rate of land conversion from state I to state C, or state M to state C) = 2.6% while δ_m_ (rate of land modification from state I to state M) = 0.96% (Gaveau et al. 2016). Under the current practises on the island, intact habitat (I) is constantly being cleared (C) and modified (M). In order to provide the economic gains of plantations and high quality timber while reducing the amount of cleared habitat, the amount of modified habitat will increase.

Considering the four possible action pathways described above (Fig. 2b), our results show that if the only aim was to increase timber production whiel reducing cleared habitat (i.e. to increase M habitat) then investment would have to be in land management (b_2_) with minimum investment in intensive restoration (b_1_) or moderate restoration (b_3_) (Supplementary figure 2). Most combinations of b_1_/b_2_/b_3_ give an increase in I, so all management actions lead to positive outcome for biodiversity (Supplementary figure 2). The magnitude of change is greater for a large budget than a small budget.

### 3. Agriculture

#### Agriculture practices and biodiversity

Loss of biodiversity and future productivity in agro-ecosystems is a major global concern. The majority of agriculture is dependent upon biodiversity, for example insects for pollination and natural pest control, which underpins a wide variety of ecological goods and services (Power 2010). Even high intensity cropping systems rely greatly on supporting and regulating ecological services and thus consideration of both production and biodiversity is essential in productive agricultural systems (Bommarco et al. 2013).

Within agricultural landscapes, land use can be divided into three main types: arable land (C), semi-natural areas such as grasslands and forests (M), and occasionally nature reserves or protected areas (I), although the latter areas are rare in agricultural setting. Semi-natural areas can range from small patches of unused vegetation such a grassy field boundaries or banks of water courses to unmanaged woody patches and set-aside fields, and the proportion of these land use types vary in different farm types.

#### Applying the decision framework: Darling Downs case study

The Darling Downs is situated 200km west of Brisbane, Queensland, Australia. The area is known for its diverse agriculture due primarily to extensive vertisol soils and humid sub- tropical to semi-arid tropic climate. The area is farmed primarily for summer and winter crops including cotton, sorghum, legumes, wheat and barley. The average amount of intact (mainly riparian vegetation), modified (disturbed semi-natural area) and cleared (mainly crop) areas within a 1km_2_ area (average farm management size) was calculated from 24 landscapes.

##### a- Objectives

The main objectives of the majority of farmers are to: 1) maximise their yield and thus income(i.e. increase C) and 2) minimize their risk from pest infestation, weeds, fire or development of insecticide resistance (Jellinek et al. 2013) (i.e. increase M or I). In addition, a third objective of maintaining biodiversity (i.e. increase M and/or maintain I) is of importance to a small but increasing number of farmers who see themselves as stewards of biodiversity and ecosystem services (Januchowski-Hartley et al. 2012; Greiner 2015) and/or understand that protecting and maintaining native vegetation will also provide private benefits (Januchowski-Hartley et al. 2012). However, in relation to the latter objective, farmers are often constrained financially. In some countries subsidies exist for certain interventions to benefit biodiversity (e.g. agri-environmental schemes in EU, US Conservation Reserve Program, Victorian Bush Tender Program) but these are voluntary and by no means global (Kleijn & Sutherland 2003; Stoneham et al. 2003; Claassen et al. 2008).

In addition, a lack of information and uncertainty of management actions that benefit biodiversity and production (e.g. Page & Bellotti 2015) discourage farmers from engaging in biodiversity friendly practises.

##### b- Actions

In order to meet these objectives while preventing any further increase in cleared habitat the main actions that farmers can take are: 1) restore cleared habitat to accommodate both agriculture and biodiversity to increase yield and farm income while minimizing the risk to biodiversity (C to M), 2) manage habitat for natural pest control on their land (C to M), 3) take land out of production and revegetate, or improve non-productive land (C to M, C to I or M to I) (Fig. 2c). This later action is rare in productive agriculture areas although agri-environmental schemes and similar programs encourage farmers to revegetate certain areas on the farm in return for a financial reward.

##### c- Costs and constraints

Generally, farms are privately owned and managed, and farmer’s budgets are often the primary constraint on the actions taken. Financial gain and risk reduction are important for farmers. In order for farmers to voluntarily revegetate parts of their farm there generally need to be both an understanding the indirect benefits biodiversity bring to production (Bommarco et al. 2013) as well as a direct financial subsidy for undertaking such action; these are available in some countries through certain agri-environment schemes.

##### d- Model

Using GIS data, we calculated average values of I=5%, M=20%, C=75%. We estimated δ_c_ (rate of land conversion from state I to state C, or state M to state C) = 7.5% while δ_m_ (rate of land modification from state I to state M) = 0.0%. Under the current management land is likely to be converted from a modified to cleared state, while intact habitats often have a certain level of protection. Conservation actions therefore need to be low budget to overcome this and prevent clearance of modified habitat (Supplementary figure 3).

We found that positive gains can be made to the amount of intact habitat for all budget sizes for all combinations of management actions when investment in land management (b_2_) ≤ 80% (Supplementary figure 3). As the budget gets larger more land can be left intact as long as investment in land management (b_2_) ≤ 90% Supplementary figure 3).

## Discussion

Creating a landscape which meets multiple objectives, such as conservation and food and fibre production, or conservation and urban amenity is not an easy task. Navigating these multiple objectives and targets while meeting other demands on natural ecosystems requires a more nuanced approach to decision making that accounts for multiple pathways towards desired system states. This is in contrast to more blunt recommendations to decision makers that emerge from orthodox dichotomies such as simply recommending a spared or shared landscape.

Focusing solely on the ‘ideal’ landscape for conservation is often unattainable and may not be an easy approach to implement by decision makers. We argue that a more useful approach is to establish incremental movement towards a landscape that better meets the goals for multiple objectives of decision makers such as biodiversity and production, supported by a transparent decision framework. Such a decision framework must include the current state of the landscape and background rates of change, a clear set of objectives, associated actions with their costs, and a total budget (Carwardine et al. 2008). Pairing this approach with a generalised system model we illustrate the potential landscape changes that can occur through different management actions in three example landscapes. The case studies demonstrate how the approach can be readily applied to a range of landscapes to inform decision making. This approach can be used by land managers or policy makers to predict the effect of different actions on the overall landscape and determine the optimum actions required in order to achieve their objectives within a given budget, and realistic timeframes.

Each of the case studies illustrates that varying the investment in different conservation actions can allow the different objectives of the landowner to be met within the available budget. This allows managers to predetermine outcomes from different actions costing different amounts, which is a valuable tool for both conservation and production.

A key benefit of our high level model over other frameworks, such as the land sparing versus land sharing dichotomy, is that it is not spatially restricted and can be used in any landscape and at any spatial scale. Each of our case studies varies in spatial scale, current landscape compositions and rate of change. We show that the current rate of habitat conversion across the landscape largely affects the outcome of conservation management emphasising the importance of including the dynamic nature of the landscape rather than a static snapshot in time. Our results consistently show that when the rate of conversion is high it takes considerably more action, and hence cost, to achieve positive conservation gains in terms of more intact habitat. Commonly the current management and rate of habitat loss is not considered in conservation management plans meaning that the cost of obtaining the desired landscape is often considerably higher than planned and thus unattainable within the available budget. The proposed system model could be used in an optimisation framework in order to provide time-dependent recommendations for investment to achieve the objective set by managers. This will allow planners and managers to plan for future targets and circumstances on the land.

The traditional land sparing versus land sharing framework focuses on selecting the landscape design to maximise the species richness of specific species of conservation concern given two alternative landscapes. It fails to consider other potential objectives or other landscape designs to meet such objectives, or the fact that the path between two landscapes may not be linear (see Path P_2_ in Fig. 1 for example). Agreeing upon objectives with stakeholders is an essential component of landscape planning (Game et al. 2013). These objectives may relate to a range of values, such as economic or social (e.g. agricultural production), in addition to conservation objectives (e.g. protection of 50% of the available land), and be incorporated into the decision making process as features to minimise or maximise or as constraints (e.g. conservation budgets or minimum food production values).

Optimising for one objective is relatively easy, however multiple objectives arise when both production and conservation demands are included often creating trade-offs and a more complex but realistic outcome.

Commonly, the cost of actions is ignored in conservation prioritisation leading to inefficient decisions and expensive mistakes (Carwardine et al. 2008). As would be expected, changes in the cost of management actions can largely affect their outcomes with greater benefits for conservation being possible when the cost of actions are low (as shown in Supplementary figures 1-3 where the increase in state I is greatest when the cost of actions are low). Failure to include the costs can result in idealistic actions being suggested that may not be financially viable. Although, in our case studies, we have made the cost of all actions the same for simplicity, different actions are likely to accrue different costs and these can be factored into the decision making process.

This research is directly applicable to land management policy. One of the criticisms of linking science outcomes to policy is that scientists ‘fail’ to ask policy relevant questions and provide an approach that is timely and consistent. The framework proposed overcomes some of these criticisms; it is generalizable, transparent and flexible. The current state of landscapes (as the starting point for the case studies) are often the result of a policy already in place. Therefore, this framework can be used as a tool where policies can be treated as hypotheses that can illustrate to decision makers the costs and trade-offs (Perrings et al. 2011; Thompson et al. 2011).

Our spatially independent multi-state landscape model provides a more realistic and feasible decision framework to aid land managers in meeting both conservation and production targets. The approach can be used globally in any landscape type so long as the objectives, actions and cost of those actions are explicitly known. This allows landscapes to be managed accordingly to meet different objectives within the available budget. This provides for more attainable and acceptable decisions to be made over the management of the landscape.

## Acknowledgements

This research was conducted with the support of funding from the Australian Research Council Centre of Excellence for Environmental Decisions (ARR and HPP) and CSIRO Julius Career Award (NAS) for the contributions to ARR. ALMC, VMA, and HPP acknowledge funding from an ARC Laureate Fellowship awarded to HPP (FL130100090).

## Supplementary Figures

**Supplementary Figure 1:**
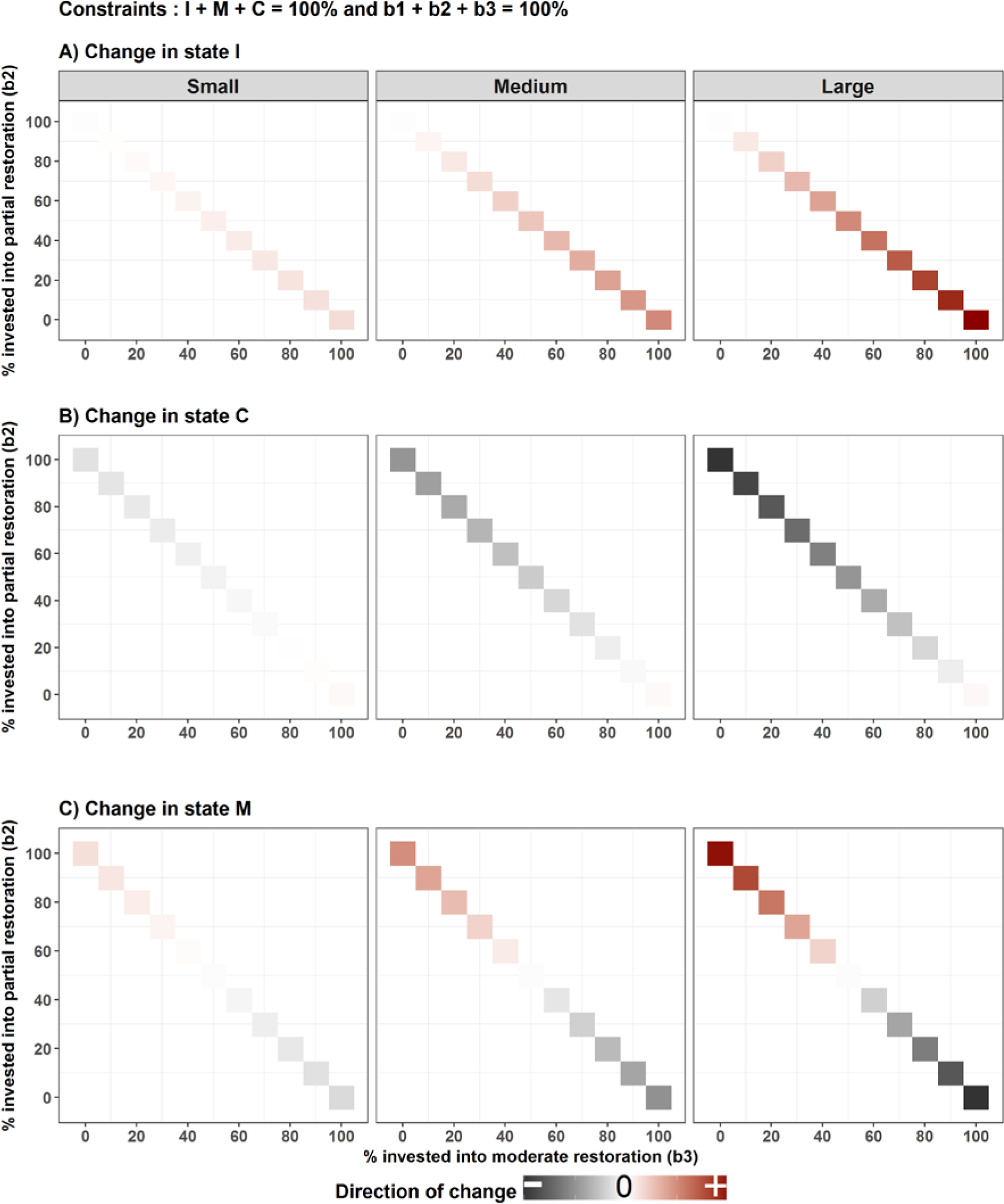
Change in the percentage of area in all three states intact (I), converted (C), modified (M) in Brisbane over one time step (e.g. one year) using our system model, when the budget is small, medium or large. This shows the possible investment in moderate restoration b_3_ (M to I; x axis) and land management b_2_ (C to M; y axis) that could lead to an increase in the amount of I area (top row), and the corresponding changes in C (middle row) and M (bottom row). Here the investment into intensive restoration b_1_, C to I, is 0%. An increase in the states is shown in red while a decrease is shown in grey; a darker colour signifies a larger change.

**Supplementary Figure 2:**
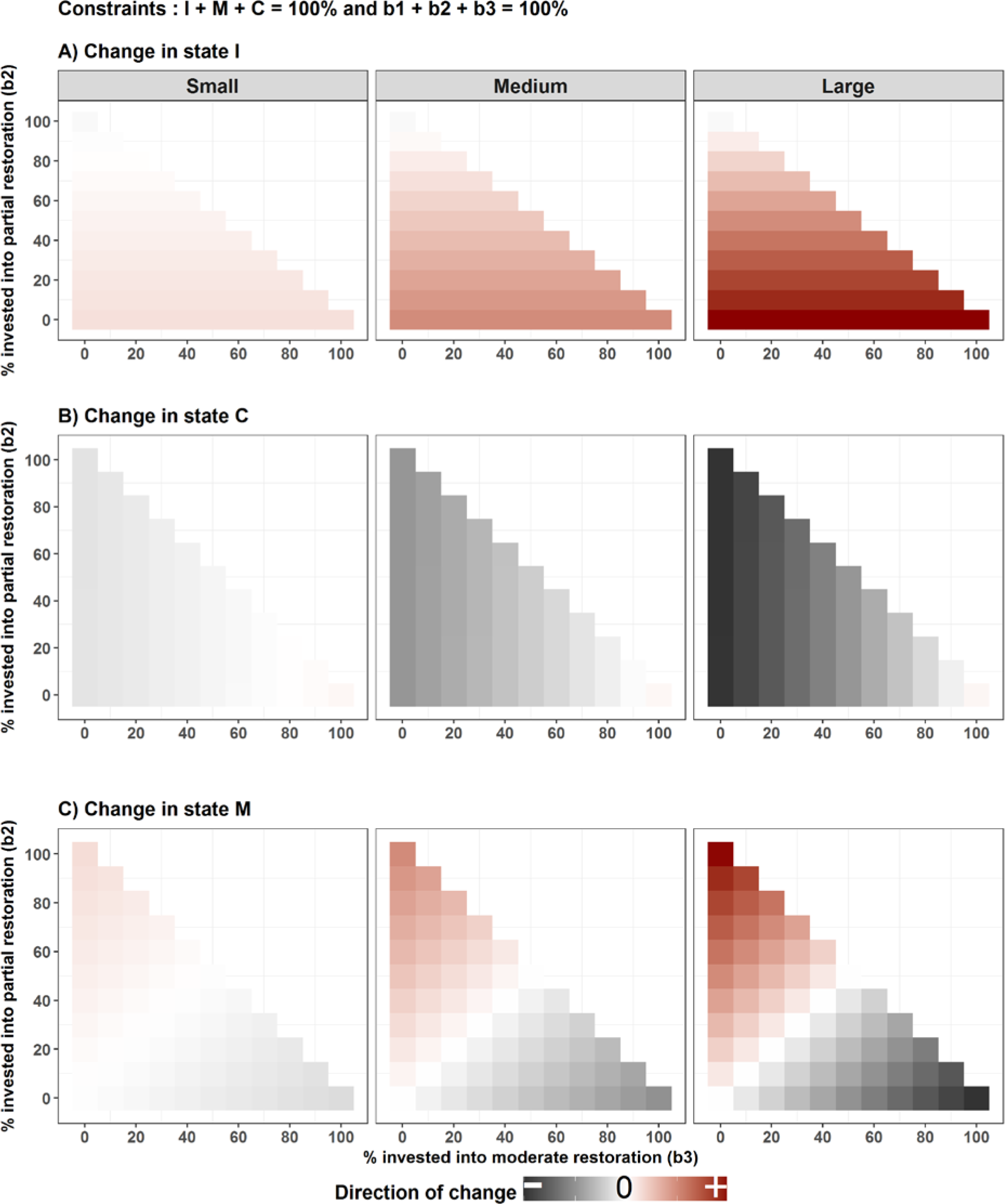
Change in the percentage of area in all three states intact (I), converted (C), modified (M) in Borneo over one time step (e.g. one year) using our system model, when the budget is small, medium or large. This shows the possible investment in moderate restoration b_3_ (M to I; x axis) and land management b_2_ (C to M; y axis), and intensive restoration (C to I; calculated as 100 – b_2_ – b_3_) that could lead to an increase in the amount of I area (top row), and the corresponding changes in C (middle row) and M (bottom row). An increase in the states is shown in red while a decrease is shown in grey; a darker colour signifies a larger change.

**Supplementary figure 3:**
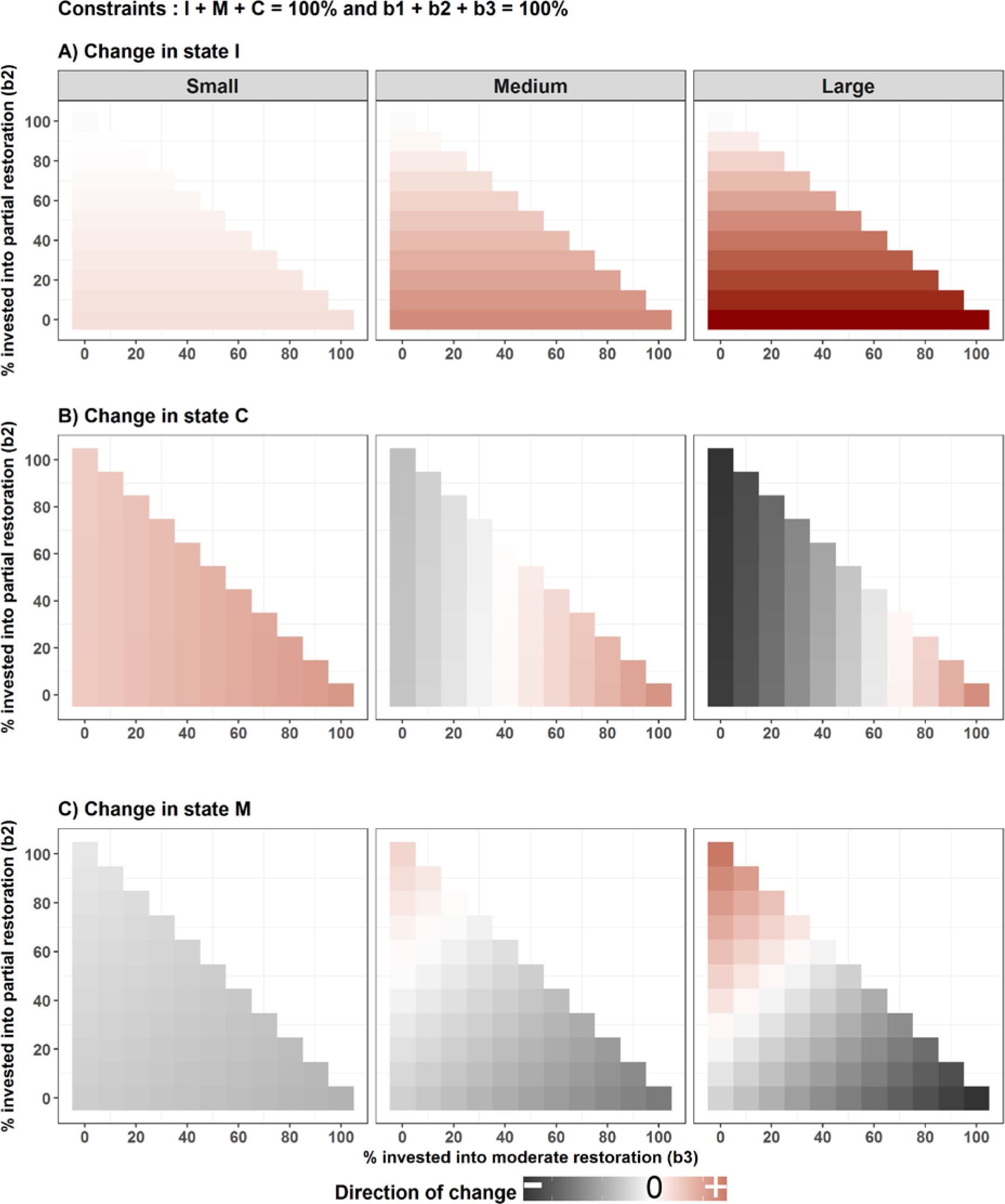
Change in the percentage of area in all three states intact (I), converted (C), modified (M) in the Darling Downs over one time step (e.g. one year) using our system model, when the budget is small, medium or large. This shows the possible investment in moderate restoration b_3_ (M to I; x axis) and land management b_2_ (C to M; y axis), and intensive restoration (C to I; calculated as 100 – b_2_ – b_3_) that could lead to an increase in the amount of I area (top row), and the corresponding changes in C (middle row) and M (bottom row). An increase in the states is shown in red while a decrease is shown in grey; a darker colour signifies a larger change.

